# Addressing technical variations in ATAC-seq data and improving motif accessibility analyses

**DOI:** 10.64898/2026.08.21.746151

**Authors:** Jiayi Wang, Emanuel Sonder, Silvia Domcke, Mark D. Robinson, Pierre-Luc Germain

**Affiliations:** Computational Neurogenomics, D-HEST, ETH Zurich, Switzerland; Department of Molecular Life Sciences, University of Zurich, Switzerland; SIB Swiss Institute of Bioinformatics, University of Zurich, Switzerland; Systems Neuroscience, D-HEST, ETH Zurich, Switzerland

**Keywords:** ATAC-seq, epigenomics, Cut&Tag, single-cell, DNA motifs, transcription factors, accessibility, differential analysis

## Abstract

Tagmentation-based methods such as ATAC-seq and Cut&Tag have provided easy ways to profile the epigenome in low-input samples and even single cells. In this contribution, we discuss forms of bias (i.e. technical variations) in tagmentation-based data, in particular ATAC-seq, and introduce three R/bioconductor packages to facilitate bulk and single-cell epigenomic data analysis, with a special focus on motif accessibility analysis. The *weightedMotifAccess* package uses weight models to enable motif accessibility analysis, including transcription factor footprint information. The *betterChromVAR* package provides a novel, analytical re-implementation of the popular *chromVAR* method that offers substantial speed improvements, eliminates stochasticity, and offers additional features. Based on this, we also propose a method, CVnorm, that outperforms alternatives in normalizing technical bias in peak count data. The computational efficiency of these tools further enables a new framework for systematically investigating synergistic and antagonistic interactions between transcription factor motifs. Finally, the *epiwraps* package streamlines the visualization, normalization, and summarization of epigenomic data.

## Background

Epigenomic data modalities, such as ATAC-seq (Assay for Transposase-Accessible Chromatin with sequencing)^[1]^ and ChIP-seq (Chromatin Immunoprecipitation followed by sequencing) or its derivatives, offer important insights into the determinants of gene expression and represent a critical complement to transcriptomics. They enable the identification of regulatory elements and their regulatory states, and can be used to infer transcription factor (TF) activity^[2]^ or attempt to reconstruct transcriptional networks, in both bulk and single-cell data ^[3,4]^. Despite the long history of methodological advances in the analysis of such epigenomic data, its day-to-day handling by end users has not enjoyed the same degree of smoothness and standardization as we have come to enjoy, through Bioconductor, for transcriptomics. One contribution in this paper is therefore a package, *epiwraps*, aimed at simplifying and streamlining some of these tasks.

The rest of the paper is aimed at ATAC-seq analysis, although the methods are largely applicable to similar assays (such as H3K27ac Cut&Tag profiles). In particular, we developed methods aimed at TF activity inference through motif accessibility, paying careful attention to the biases present in ATAC-seq data.

### Technical biases in ATAC-seq data

As we see it, there are three inter-related types of technical biases or variations in ATAC-seq data, which we call enrichment bias, fragment length bias, and GC bias. These biases are especially important when their magnitude differs across samples. We illustrate this in Figure 1 with ATAC-seq profiles of the same sample, created using variations of the protocol^[5]^, but emphasize that similar variations (usually milder) regularly occur even when using the same protocol.

**Figure 1:**
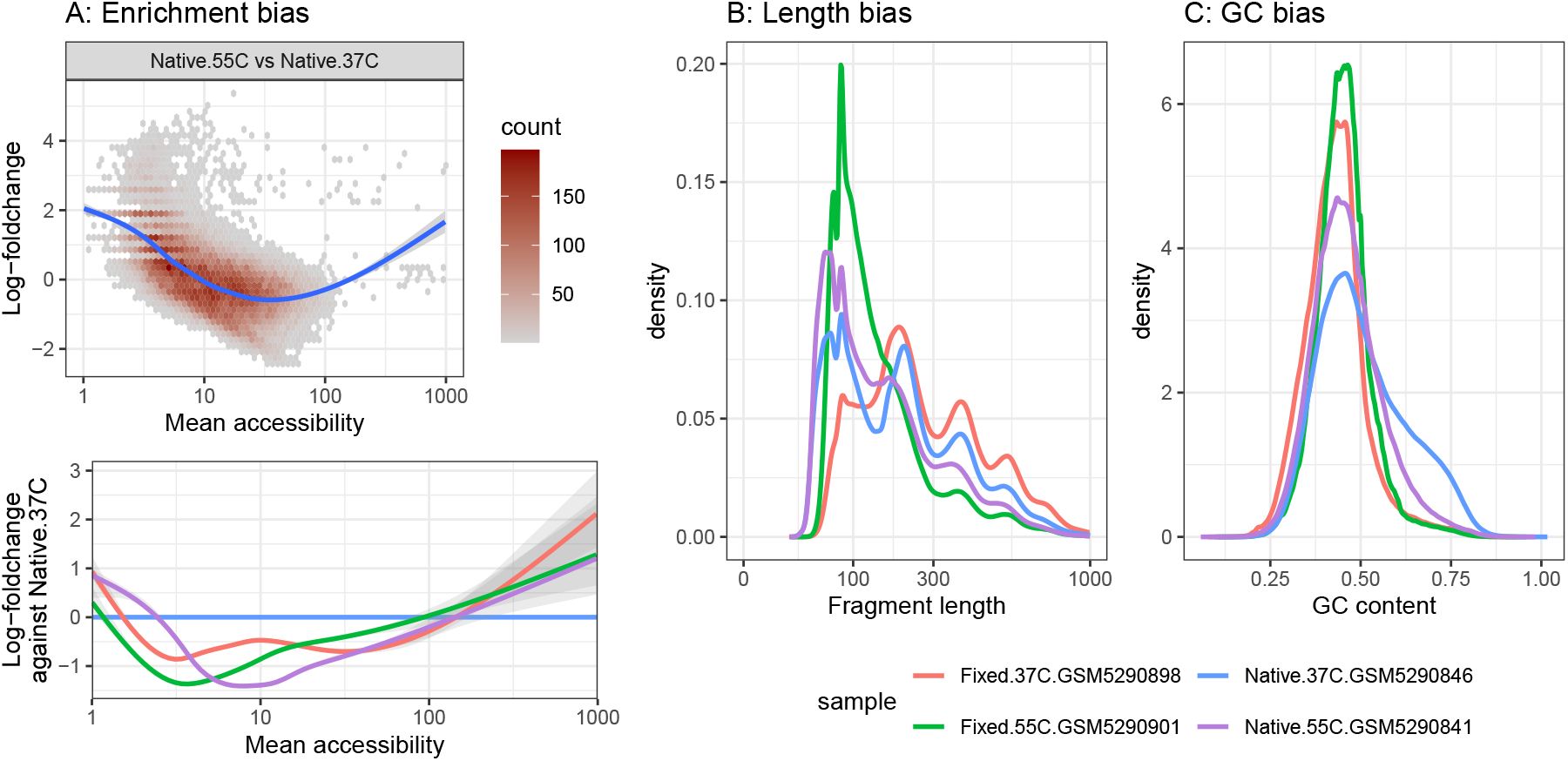
Illustration of variations in ATAC-seq biases, using the Zhang et al. dataset ^[**5**]^. **A:** Enrichment bias is when a sample shows a higher concentration of the reads in high-accessibility regions. The example represents a very strong difference in the degree of enrichment due to ATAC-seq performed at different temperatures. It can be visualized for instance either as a slope/curve on the minus–average (MA) plot. Lines represent Generalized Additive Models. **B-C:** Differences in the distributions of fragments’ length (**B**) and GC content (**C**) across four profiles of the same sample. Importantly, while we use different protocols to highlight these variations, they also very often arise in samples prepared with the same protocol.

Enrichment bias has long been recognized in related ChIP-seq data^[6]^, and represents the degree to which the reads are concentrated in high-signal regions. It can for instance be visualized in the traditional minus–average (MA) plot as a sloped (or curved), rather than a horizontal, trend line (Figure 1A, top), or as an enrichment over the mean (Figure 1A, bottom), which is not always monotonous. Fragment length bias represents the relative abundance of fragments of different sizes. A hallmark of ATAC-seq data is its periodic distribution of fragment sizes into nucleosome-free, mono-nucleosome containing, and poly-nucleosome containing fragments, but their relative abundance often varies across samples due to differences in tagmentation efficiency (Figure 1B). Finally, GC bias refers to the relative prevalence of guanines and cytosines in the fragments amplified and sequenced. ATAC-seq fragments generally have slightly higher GC content than would be expected by chance based on the genome composition, which has a number of potential explanations. The first is the cleavage bias of the Tn5 transposase itself, which was shown early on to have a higher tendency to cut at sites presenting such nucleotides^[1]^, even in naked DNA^[7]^. While correcting for this bias is critical for certain tasks such as footprint identification^[7,8]^, for many other tasks, such as differential analysis across conditions, it is problematic only if the magnitude of the bias varies across samples (as illustrated in Figure 1C). Such variations can plausibly arise if the amount of Tn5 (relative to cell concentration), its efficiency (e.g. through temperature), or the duration of the incubation changes. Another often cited explanation for GC bias is PCR amplification^[9,10]^. However, in most contemporary ATAC-seq protocols, the GC bias of PCR amplification is minimal (except in extreme cases, e.g. long non-interspersed GC stretches^[11]^). To see this, we can investigate PCR duplication events in single-cell ATAC-seq, which are the same fragment sequenced multiple times with the same unique molecular identifier (UMI) and cell barcode. When we then compare the GC content of duplicated, versus non-duplicated fragments, we do observe a slight shift towards higher GC content (Supplementary Figure S1A), however we would argue this is an indirect effect. Indeed, the probability of a successful PCR decreases with the length of the fragment^[12,13,14]^, such that duplicates tend to come from shorter fragments (Supplementary Figure S1B); since shorter fragments, which are nucleosome-free, typically come from GC-rich regulatory elements, they show a GC enrichment (Supplementary Figure S1C). The GC bias observed in duplicated fragments is therefore accountable by their length bias (Supplementary Figure S1C-D). That said, there remains differences in GC content between certain ATAC profiles that are neither due PCR bias, nor reducible to fragment length (Figure 1B-C).

The three types of biases are strongly intertwined because highly-accessible regions tend to be GC-rich, nucleosome-free regions. As such, a bias in one of these dimensions is likely to create secondary bias in the others, although they are not entirely coupled either. Indeed, an initial difference in, for example, tagmentation efficiency, can bias the results in multiple, competing ways. On the one hand, as the number of insertions in a genome increases, the length of the fragments between them could be expected to decrease. On the other hand, since highly-accessible sites with high affinity for the Tn5 have already had an insertion, further insertions events are more likely to occur in less insertion-prone regions, thus leading for instance to a lower enrichment for GC-rich, nucleosome-free regulatory elements. This is for instance observed in native chromatin (Figure 1A, bottom^[5]^), however the opposite is observed in fixed cells^[5]^, suggesting a multifaceted impact of the protocol. Either way, further differences introduced by the length bias of PCR amplification, as well as the sequencing platform^[13]^, can then compound such initial biases.

In most cases, such variations across samples are technical. Indeed, even when comparing across very different cell types in the same single-cell ATAC-seq sample, one observes no variation in the fragment length distribution, and very limited differences in GC content (Supplementary Figure S1E). It is therefore advisable to correct for such variations for most downstream analysis, and hence methods that account for such bias typically outperform alternatives in downstream tasks^[10,2]^. A particularly successful method in the context of TF activity inference is *chromVAR*^[9]^, which was developed for single-cell ATAC-seq data, but was also shown to be very powerful for bulk ATAC-seq data^[2]^. Its success can be attributed in good part to its attention to enrichment and GC bias. Briefly put, *chromVAR* uses peaks with a similar average accessibility and a similar GC content as background to correct for sample-specific GC and enrichment biases. It does not, however, pay explicit attention to length bias. Furthermore, its permutation-based approach does not scale to large single-cell datasets, and introduces an undesirable element of stochasticity. Attempts have been made to develop a faster re-implementation of the method, still within a permutation framework^[15]^, but scalability has remained an issue. Here, we therefore developed alternatives to this approach, and expanded their applications.

## Results

### ATAC-seq bias correction through weighted counts

Recognizing the technical biases caused by GC content and fragment lengths (FLs) in ATAC-seq, we developed a weighting framework that assigns weights to both fragments and accessible regions (peaks), thereby improving peak- and motif-level differential accessibility analysis without the need for background permutations. Briefly, fragments are stratified by GC content and fragment length bins and the bins are assigned sample-specific weights to harmonize their relative frequency across samples. This is followed by cyclic loess normalization on the peak-level weighted fragment counts (similarly to *csaw* ^[6]^ for ChIP-seq) to correct enrichment bias (Supplementary Figure S2, see Methods). The weighted count matrix can be generated using the `getWeightedCounts` function in the *weightedMotifAccess* package and can be used for downstream peak-level differential analysis, as well as for motif accessibility. To assess whether our weight model mitigates technical biases, we revisited the difference between ATAC-seq profiles generated at 55 ^*°*^C vs. 37 ^*°*^C (as in Figure 1A). The original counts exhibit a systematic bias, with the fitted trend deviating from zero across average accessibility levels. After applying the weight model, the trend is largely centered around zero, with a more symmetric distribution (Figure 2A).

**Figure 2:**
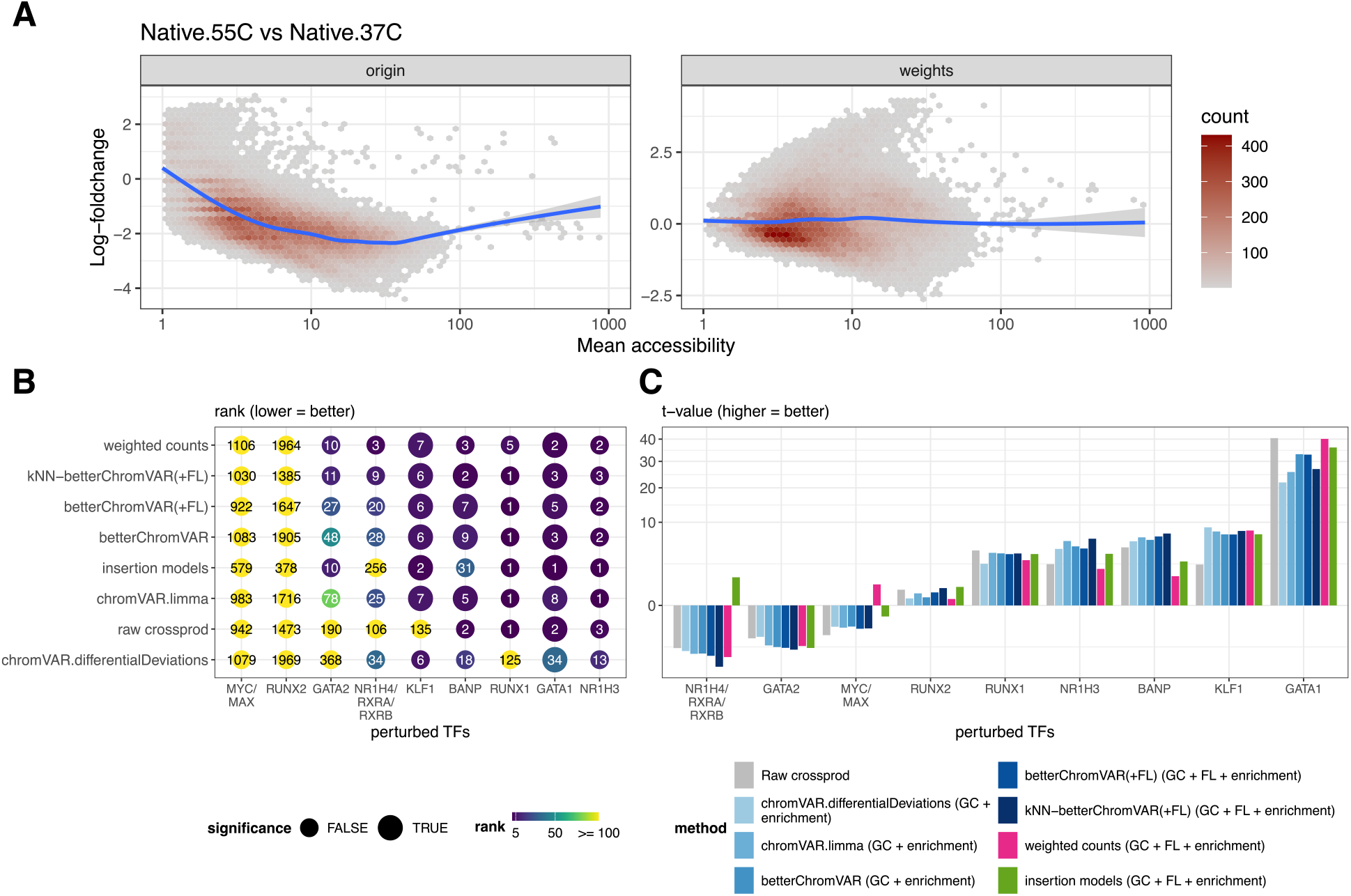
**A**: MA plots comparing two samples performed at different temperatures, using original and weighted counts from^[5]^. Blue lines represent Generalized Additive Models. **B-C**: Benchmark of the different method variants on the recovery of differentially active TFs, based on Gerbaldo, Sonder et al.^[2]^. The number and color in **B** indicate the rank of the perturbed TF (the top-ranked TF if multiple). Significance is defined as an adjusted p-value < 0.05. Ordering of methods and datasets in panel **B** is based on the mean square root of ranks; datasets in panel **C** are ordered by the mean t-value. Parentheses indicate the biases corrected by each method. The *chromVAR* values were averaged over 2 random seeds. Raw crossprod indicates the direct application of *limma-voom* to the cross-product of the original peak-level count matrix and motif matches. The NR1H4 and NR1H3 perturbations are ligand-based activation, while all other cases are downregulation of the TF (using a degron for BANP, and CRISPRi KD for the others). The sign of the *t*-values in **C** was therefore inverted so that a positive value is expected in all cases. For MYC and NR1H4, their obligatory binding partners (MAX and RXRA/B, respectively) were detected and therefore included in the set of perturbed TF motifs for evaluation.

Next, we investigated whether the weight model improves differential motif accessibility analysis using the same benchmark datasets as in the previous evaluation^[2]^. In each dataset, a single TF is perturbed, and an effective method should prioritize the corresponding motif as the most significant, i.e. top rank and high absolute t-statistics. Overall, methods correcting for GC, fragment length (FL), and enrichment biases tend to rank among the top-performing approaches. The weight model compared favorably with chromVAR-based methods, assigning relatively low ranks to all perturbed TFs except MYC and RUNX2 (Figure 2B). Consistent with recent benchmark results, all methods failed to identify RUNX2 and MYC, which also failed to show significant peak-level differences, suggesting the absence of a clear treatment effect^[2]^.

Taken together, these results suggest that the proposed weighting framework effectively reduces technical biases in ATAC-seq data and improves the accuracy of downstream differential motif accessibility analyses. In addition, the weighted counts could be used for peak-level differential accessibility analysis.

### betterChromVAR: fast, analytic motif deviations

The *chromVAR* method^[9]^ is highly sensitive^[2]^ and widely used, but limited by the imprecision and computational costs of running background permutations. We therefore developed an analytical re-implementation of the method (see the *betterChromVAR* package), which entirely bypasses the permutations (thus making it deterministic), and achieves considerable speedup by doing most of the computation in the space of bias bins, rather than in the peak space (see the Methods section). The method is substantially faster than the original *chromVAR* and the C++ re-implementation included in the *ArchR* package^[15]^, and also operates within a small memory footprint (Figure 3A-B).

**Figure 3:**
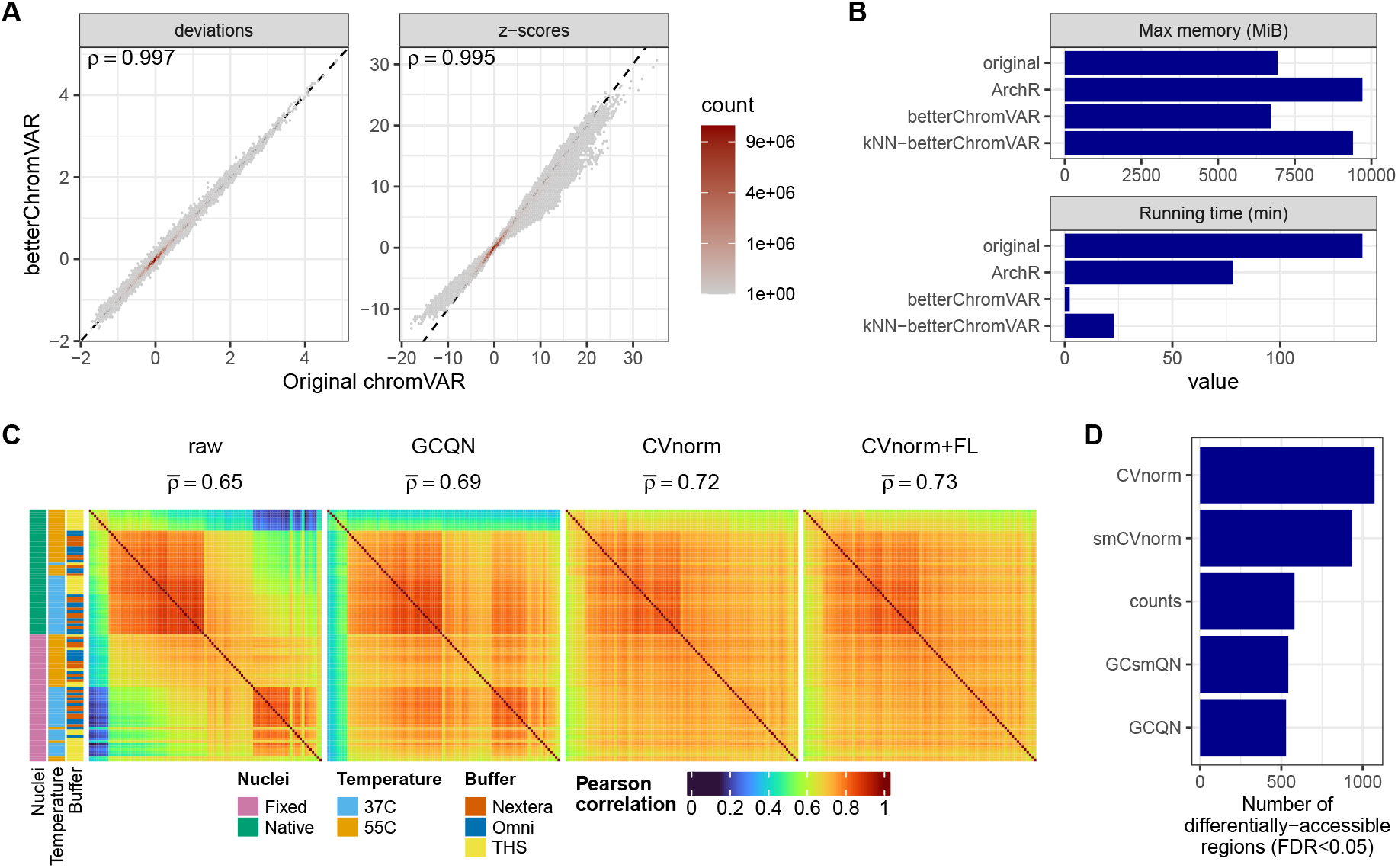
**A-B:** Comparison of *betterChromVAR* to the original *chromVAR* on a dataset of 30839 cells, 157811 features and 2065 motifs (non-neuronal cells from Waag *et al*. [19]), using 100 background iterations. **A:** Comparison of the per motif/sample deviations (left) and z-scores (right). (The *chromVAR* values were averaged over 2 random seeds.) **B:** Memory usage (top) and running time (bottom, 1 CPU) of the two implementations, as well as the C++ implementation from *ArchR*. The time needed to scan for motif matches was subtracted from all methods. **C:** Pairwise correlation of log1p(counts) between ATAC-seq profiles of the same sample, produced using different reagents and temperatures^[5]^, and normalized using quantile normalization in GC bins (GCQN) or using CVnorm, the latter with classical chromVAR biases or supplemented with fragment length (FL) bias. Strong correlation patterns seen in the uncorrected data are strongly corrected by CVnorm. **D:** Usage of CVnorm in a bulk ATAC-seq dataset^[19]^ leads to the discovery of more differentially-accessible regions than alternatives. smCVnorm and GCsmQN respectively indicate CVnorm and GCQN with smoothing (based on the acute stress group).

We next sought to further improve the method. First, we enabled it to make use of non-binary annotations, such as probabilities obtained by binding prediction models^[16,17]^. Second, to reduce bias related to cell type abundance^[18]^, we enabled a more straightforward balancing of expectations by giving the same weight to the different cell types. Third, to make bias estimation at the single-cell level more robust, we implemented an empirical Bayes shrinkage approach across cells of the same group (see Methods). We did not however find convincing evidence of this shrinkage improving cell-level inferences (data not shown). Finally, we integrated the third type of bias, namely fragment length bias, into the generation of the background bins. To this end, the `peakCountsFromBAM` and `peakCountsFromFrags` functions of *epiwraps* compile, while counting overlaps, the median fragment length per region.

*betterChromVAR* produced motif deviations strongly correlated with the original *chromVAR* (Figure 3A and that led to comparable (similar or slightly better) TF recovery (Figure 2B-C).

### betterChromVAR with kNN-based background

The *scPRINTER* package, an expansion and scalable version of the *PRINT* method ^[16]^ developed by Ruochi Zhang in the Buenrostro lab, includes a reimplementation of *chromVAR* with an important modification. First, rather than cutting the bias space into bins, a continuous, multidimensional bias space space is created from which *k* nearest neighbors (kNN) are selected as background. This approach scales nicely to the inclusion of more dimensions of bias. The mean expectation and its variance are then computed across the *k* neighbors, analogously to the classical *chromVAR* permutations, although sped up by a GPU implementation.

We reasoned that such an approach could also benefit from replacing the expensive permutations with analytical estimates. We implemented such an approach (in the `getBackgroundKNN` and `computeDeviationsFromKNN` functions of *betterChromVAR*), with two additional features. First, computing kNNs across the multidimensional bias space makes the assumption that each bias dimension has an equal importance, which is not necessarily the case. To address this, we scale the dimensions by the proportion of the variance in peak over-dispersion they explain. Second, *chromVAR* backgrounds have the drawback that, for some motifs, a fairly large proportion of the background peaks actually harbor the motif. We took advantage of the way we implemented the kNN-based approach to have the option of masking those peaks from the background in computing motif expectations (see methods). The analytical kNN variant of *betterChromVAR* is still much faster than the original *chromVAR*, but slower and requires more memory (Figure 3) than the bin-based analytical approach (more memory can be used to linearly increase speed). Masking motif-harboring background peaks led to worse results, and we thus disable it by default (Supplementary Figure S3). Instead, the weighing of the bias dimensions slightly improved results (Supplementary Figure S3). The best kNN variant showed small improvements over the bin-based variant in TF recovery (Figure 2). At the single-cell level, however, it resulted in motif deviations that showed slightly lower intra-replicate correlations and slightly higher within-class variance (Supplementary Figure S4).

### Motif deviations using insertion models

Some transcription factors (TFs) leave characteristic footprints, i.e. patterns of Tn5 insertion sites around their binding sites, which have been used in some differential analysis approaches^[20,21]^, but are not currently used in *chromVAR*-like approaches. We therefore developed an insertion model to incorporate these patterns into a chromVAR-like workflow. Our strategy is to give more weight to insertions that match the expected insertion profile around the motif of each TF. The method first computes an average insertion profile around each motif, and uses it to weigh insertion events according to this expectation. The weighted counts are then aggregated across motif matching peaks (Supplementary Figure S5, see Methods) and can be generated using `weightedInsertions` function in *weightedMotifAccess* package. Those aggregated weighted counts are then compared against similar background regions using `computeDeviationsWeighted` function in *betterChromVAR* package. The insertion model performed generally well across datasets, achieving improvements over standard *chromVAR* in TFs that have a clear footprint (GATA1/2 and KLF1), while performing more poorly on BANP and NR1H4 (which has a poor ATAC footprint). Notably, however, the insertion model was the only method to recover the correct direction of effect for NR1H4.

### Normalizing out technical biases with CVnorm

Methods that correct for GC bias, such as smooth quantile normalization in GC bins^[10]^, have been shown useful not only for motif analysis, but also for peak-level differential accessibility analysis^[10]^. We therefore reasoned that a normalization based on the (analytic version of the) *chromVAR* strategy could achieve similar results, and additionally correct for other biases, by adjusting samples’ peak accessibility based on the background bias. We implemented this as `CVnorm` (in the *betterChromVAR* package) and, inspired by smooth quantile normalization^[22,10]^, additionally included an optional shrinkage of the bias correction when the bias appears to be explained by experimental groups (or other grouping provided), denoted here as smCVnorm.

To evaluate whether the method successfully corrects technical bias, we used the same dataset as in Figure 1, where (relatively shallow) ATAC-seq profiles of the same sample were generated using variations of the protocol ^[5]^. In the uncorrected data, strong patterns of pairwise correlation are seen, which are partly corrected by quantile normalization in GC bins (GCQN), but more accurately corrected by CVnorm (Figure 3C). CVnorm additionally using fragment length bias further improves the correction, albeit more modestly, in the set of samples that were least correlated to the others (Figure 3C). We note, however, that not all differences between protocols could be corrected in this fashion.

Finally, to test that biological signal is preserved through this correction, we next turned to a bulk ATAC-seq dataset from one of our recent studies about the habituation to stress in the mouse hippocampus^[19]^, and checked how many differentially accessible regions (DARs) were identified when using each bias normalization method. CVnorm retrieved the largest number of DARs with FDR <0.05 (Figure 3D), and the DARs unique to CVnorm were strongly enriched for the glucocorticoid receptor motif (adjusted p=8.9e-33), as expected from such a stress paradigm.

### Motif interaction analysis

The high computing speed of *betterChromVAR* enables a more systematic investigation of interactions (i.e. synergies or antagonisms) between motifs. We developed a simple framework to help such analyses, and illustrate it with the classical example of the activation of the glucocorticoid receptor (GCR), encoded by the NR3C1 gene, using ATAC-seq data from^[23]^. We first identified (using the `discoverMotifInteractions` function of *weightedMotifAccess*) motifs with a significant interaction with GCR in terms of the accessibility response (Figure 4A-B). We then explored these in more detail, using the `exploreMotifInteraction` function (Figure 4C). Specifically, the function uses *betterChromVAR* to analyze the adjusted (and set-size-normalized) deviations of regions harboring either motif of a pair, or their combination at specific distance ranges. In the case of a synergistic binding, sites with the two motifs should show larger deviations than sites with either motif alone, and this interaction should disappear when the distance between the motifs is too large. This is exactly what one observes for known interactors RELB and AP-1 members of the JUN/FOS family, but also for some other TFs not known to interact with GCR (Figure 4C).

**Figure 4:**
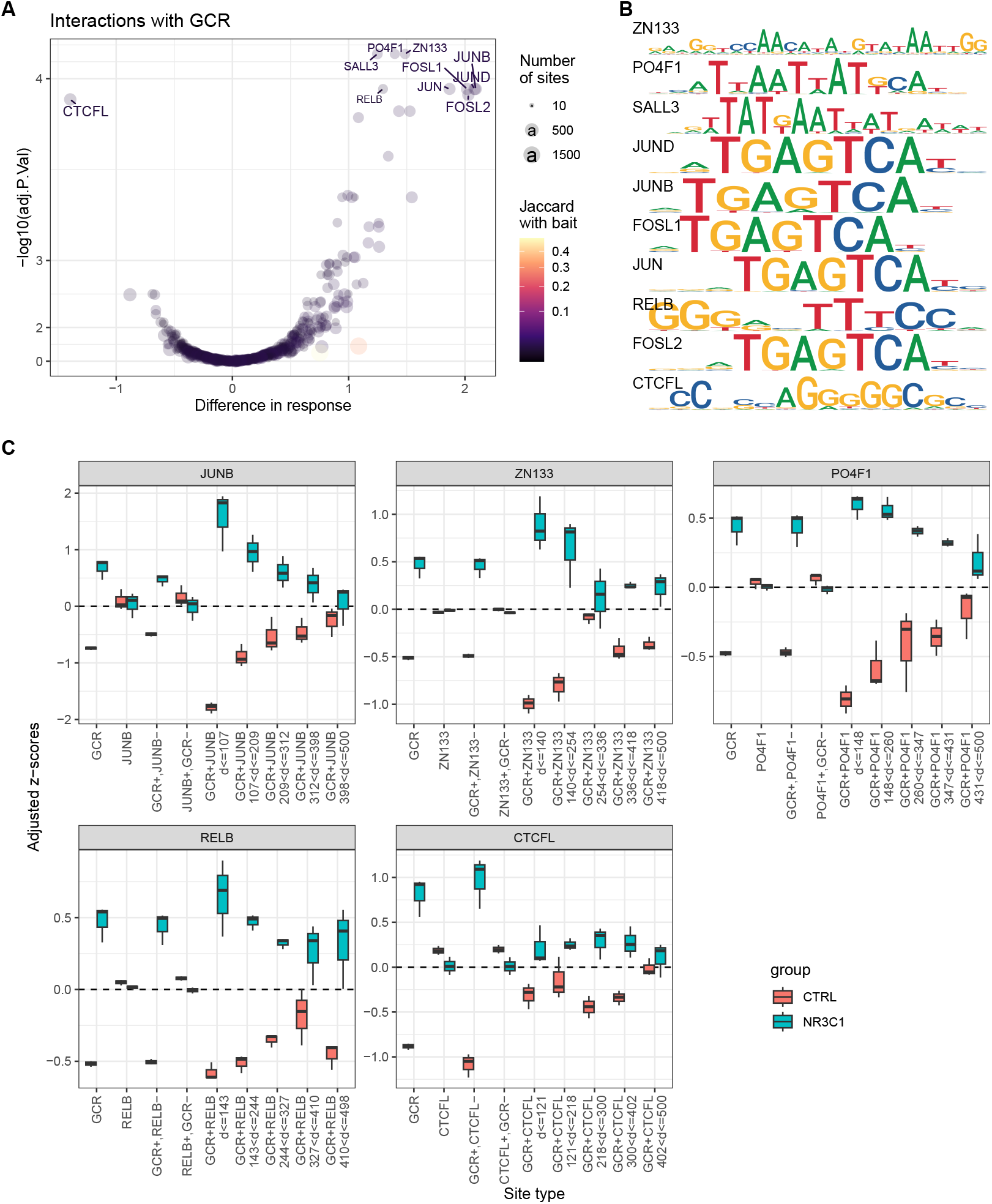
Motif interacting with GCR. **A:** Motifs that modulate GCR activation. Plotted are the difference in response (and significance thereof) of sites harboring both a GCR motif and the motif plotted, in comparison to all sites harboring a GR motif. These are the results of the `discoverMotifInteractions` function. **B:** Top motifs identified as candidates GCR-interacting motifs include the well-known interactions with JUN-family members (and AP-1 more generally), as well as that with RELB, known to form a complex with GCR. **C:** Further investigation of the top candidates using the `exploreMotifInteraction` function. Deviations are computed for sites that harbor either of the motif or their combination, the latter split into quantile distance bins (ranging from 0 to 500bp) to capture interactions decaying over distance. As expected, bona fide interactions (such as with JUN/JUNB or RELB) show stronger deviations when the two motifs are present, decaying when the motifs become too distance from each other.

### Streamlined epigenomic data visualization with *epiwraps*

Built on core Bioconductor infrastructure, the *epiwraps* package is meant to simplify summarization, normalization and plotting of epigenomic data. It was initially developed in a teaching context, to lower the barrier to entry for researchers without an extensive bioinformatics background by wrapping complex workflows into intuitive functions. The package is designed around three main principles: 1) the separation of data processing and reading from plotting, and leveraging Bioconductor data structures to enable interactivity and flexibility; 2) a more systematic approach to normalization, enabling the use of similar normalization methods across tasks (e.g. visualization and differential analysis); and, 3) a simple, intuitive interface.

The package includes, among other things:

- A flexible function (bam2bw) for summarizing alignments into bigwig files, especially useful to distinguish different ATAC-seq signals (e.g. nucleosome centers vs TF binding sites and footprints, see e.g. Figure 5A). Binning is performed directly on the run-length encoding for increased speed.
- Popular epigenomics normalization methods, including background (or SES) normalization^[24]^, common peak normalization (also known as MA-norm ^[25]^), top or enrichment normalization, and S3norm^[26]^. Importantly, the same normalization method can be applied to both read counts and coverage tracks, enabling consistency between, for instance, differential analysis and visualization.
- A wrapper for visualizing signal tracks in a single region (genome-browser), seamlessly accepting a variety of inputs, and including for instance an automatic combination of coverage track and heatmap to summarize replicates (Figure 5B).
- A streamlined interface for visualizing signals at multiple regions (e.g. Figure 5A), splitting data import from visualization for greater flexibility. Specifically, the function `signal2Matrix` reads the signal around sets of genomic regions, and stores it in a `EnrichmentSE` object (inheriting from Bioconductor’s `SummarizedExperiment`) which enables easy manipulation (e.g. subsetting/reordering), annotation, as well as post-processing (e.g. scaling/normalization) and downstream analysis. Finally, the `plotEnrichedHeatmaps` wraps around the *EnrichedHeatmap* package^[27]^ to plot heatmaps directly from the `EnrichmentSE` objects.
- A number of functions to facilitate quality control, clustering, and visualizing region overlaps.

**Figure 5:**
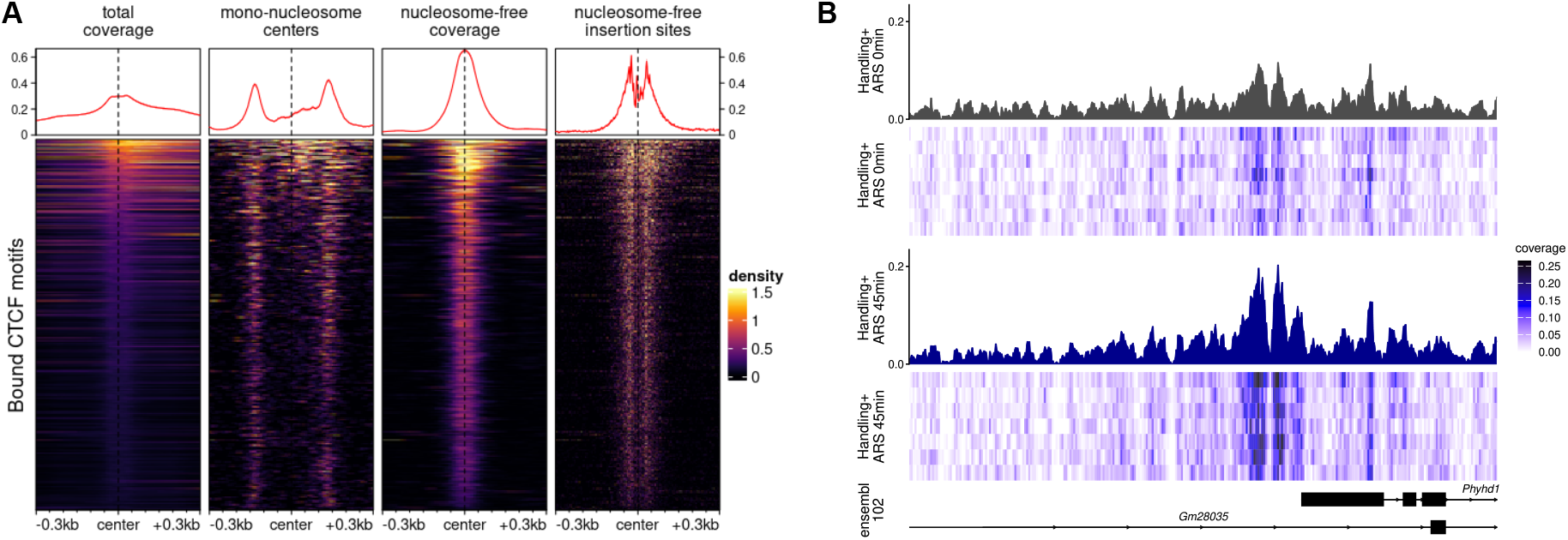
**A:** Illustration of flexible bigwig file generation and plotting using *epiwraps*. Subsets of the signal from the same T-cell ATAC-seq profile are shown around bound CTCF motifs. **B:** Example representation of single regions, offering a combination of the classical signal with heatmaps to better visualize variability across samples (`ggSignalTracks`). The data is from the mouse hippocampus after acute restraint stress (ARS)^[19]^.

## Conclusion

This contribution presents tools to facilitate epigenomic data analysis. In particular, we explored three interconnected classes of technical variations (enrichment, fragment length, and GC content biases) particularly relevant for tagmentation-based data, and methods to address them. For bulk data, we propose a simple normalization method that accurately addresses all three biases, successfully eliminates most technical sources of variation, and increases the power of differential accessibility analysis.

We then turned to motif accessibility analysis, and developed a weight-based strategy that addresses bias directly at the quantification stage. While highly successful in bulk data, the method is not scalable to single-cell analyses due to its reliance on loess fits. To this end, we therefore explored another avenue with *betterChromVAR*. By replacing the traditional permutation-based approach of the popular *chromVAR* method with an analytical framework, *betterChromVAR* makes motif deviation analysis not only deterministic but also scalable to contemporary large single-cell datasets. We further extended the analytic approach to the kNN-based background alternative used by *scPRINTER* ^[16]^. Within the resolution of our benchmark, the best kNN variant (including fragment-length bias) appeared to be slightly better at recovering perturbed TFs, but led to slightly more variable estimates at the single-cell level and took considerably longer to compute. We therefore recommend as a highly-scalable default the bin-based approach unless working with bulk data with fragment length bias. Finally, in conjunction with the *weightedMotifAccess* package, *betterChromVAR* can additionally include TF footprint information.

Beyond saving time and computational resources, *betterChromVAR*’s massive gain in computational efficiency enables systematic combinatorial analyses. We illustrate this through a new framework for studying distance-dependent synergies (or antagonisms) between TFs, as illustrated with the glucocorticoid receptor and FOS/JUN TFs.

The same set of methods could be used replacing raw motif matches with more refined binding probabilities^[16,17]^, although we did not explicitly study this here. In addition, while the methods presented in this manuscript (*epiwraps* excepted) focus on ATAC-seq analysis, we believe they are very likely to be applicable to similar epigenomic assays, such as Cut&Tag profiles. However, further research will be required to establish their performance and utility across these related data modalities.

In conclusion, the software suite presented here balances statistical rigor with computational economy, providing the epigenomics community with a highly scalable, deterministic, and user-friendly toolkit to decode TF dynamics.

## Methods

### betterChromVAR

The analytic implementation of *chromVAR* replaces stochastic background sampling with an analytical framework and computes expectations in the space of bias bins.

### Backgrounds bins

The enrichment- and GC-biases of ATAC-seq ATAC-seq are correlated, as high-accessibility regulatory elements tend to be GC-rich. To decorrelate these features of peaks, *chromVAR* uses a Mahalanobis transformation. The transformed 2D space is then discretized into a regular grid *B* of *bs* × *bs* bins (default `bs=50`). Each peak is assigned to a bin using its coordinates in the transformed space. Up to this point, this is identical to the original chromVAR, although we include the option to add a third bias dimension, namely median fragment length, in the creation of the background bins.

### Bin-to-Bin Probability Matrix (P)

In the traditional *chromVAR*, background peaks are randomly sampled from the same bin (or nearest bins if it is too sparsely populated). The similarity between bins *u* and *v* is defined by a Gaussian kernel with bandwidth *w*:

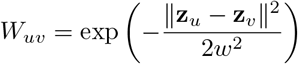

where ∥**z**_*u*_ − **z**_*v*_∥ is the Euclidean distance between bin centers. The probability of selecting a peak *q* in bin *v* as a background match for a peak *p* in bin *u* is therefore defined as:

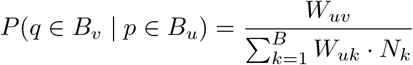

where *N*_*k*_ is the number of peaks assigned to bin *k*. Because this probability is identical for all peaks within bin *v*, we define the bin-to-bin transition matrix 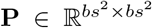 where **P**_*uv*_ represents the probability of selecting *any* single peak from bin *v* given a starting peak in bin *u*.

### Analytical Expectation and Variance

Let *C* be the matrix of observed fragment counts in peaks across samples *S*. We first aggregate *C* into a bin-count matrix *K* (*bs*^2^ ×|*S*|), where *K*_*us*_ is the sum of counts in bin *u* for sample *s*. For each bin in each sample, a background expectation can be computed from the summed counts of peaks multiplied by their probability of being selected as background. The expected count *E*_*us*_ and variance *V*_*us*_ for a peak belonging to bin *u* in sample *s* are therefore calculated as:

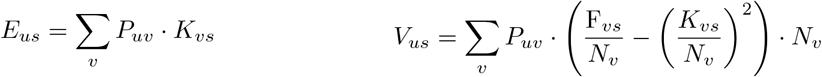

where *v* iterates over bins, F_*vs*_ is the sum of squared counts for peaks in bin *v* for sample *s*.

### Motif-Level Aggregation

Let *ϵ*_*p*_ be the global expected fraction of reads for peak *p* (e.g. calculated as the mean across cells or groups). For a motif *m*, let *A*_*m*_ be the set of peaks that contain the motif. The global expectation for motif *m* in sample *s* is:

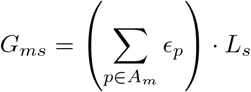

where *L*_*s*_ is the total library size (total counts in peaks) for sample *s*.

The total observed accessibility of the motif in sample *s* is 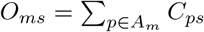. The background expectation *µ*_*ms*_ and standard deviation *σ*_*ms*_ for the motif are aggregated from the peak-level expectations:

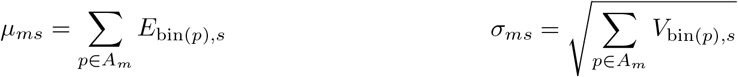

The last steps are similar to the original *chromVAR*: the final bias-corrected deviations and z-scores are computed as:

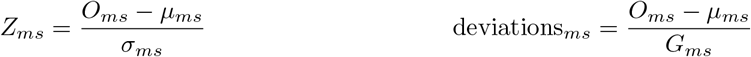

In practice, all of this is implemented as matrix operations for computational efficiency.

### Empirical Bayes shrinkage of the bias matrix

In sparse and noisy single-cell data, the background expectation at the single-cell level can be noisy. We therefore implemented optional shrinkage to the observed bin frequencies. The observed proportions of total counts found in each bin for a given sample are shrunk toward a prior which can either be defined as the average for the same bin across cells (or cells of the same type/grouping), or as a 2D-smoothed version of the same cell’s bin proportion matrix. In both cases, the shrinkage is done using an Empirical Bayes (method of moment) approach, i.e. the posterior counts of bin *u* in sample *s* is estimated as:

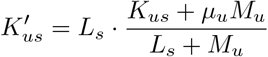

where *L*_*s*_ is the total counts for sample *s, µ*_*u*_ is the prior bin proportion, and *M*_*u*_ is the bin-specific precision parameter estimated from the observed weighted variance of the bin across samples.

### betterChromVAR with kNN-based continuous background

The `getContinuousBackground` function first constructs a peaks × *bias* matrix and decorrelates the bias dimensions as done in *chromVAR* or *betterChromVAR*, with this difference that it accepts an arbitrary number of dimensions. To make multidimensional distances more meaningful, we then scale the bias dimensions using importance weights *w* that are calculated by how much of the over-dispersion (as a proxy to inter-sample variability) is explained by each dimension. Specifically, we fit the over-dispersion 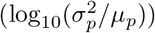 on a second degree polynomial (i.e. *y*(*x*) = *x* + *x*^2^) for each each dimension, and use the *R*^2^ of the model as weight. The transformed bias dimension is then multiplied by 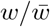. Using this weighted continuous bias space, we compute the top *k* nearest neighbors as background for each peak, as done in *scPRINTER* ^[16]^ (except using *BiocNeighbors*).

Let **W** ∈ {0, 1}^*M ×N*^ denote the sparse binary matrix indicating the presence of motif *m* in peak *p* (or an equivalent with probabilities between 0 and 1), and 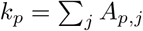 be the total neighborhood degree of peak *p*. Cell-level background expectations and analytical variances are mapped directly from the peak count matrix **X** via a single sparse projection:

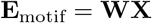

The variance *V* is then computed as the expected value of the squares minus the square of the expected values:

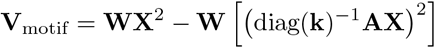

The second term, however, involves the creating of a dense cells × peaks matrix which, for large single-cell datasets, would be prohibitive to store in memory. This part is therefore run in chunks. Finally, the cell-level deviations and z-scores are computed as for other methods.

### Optional soft masking of the foreground motif from the background

An issue with *chromVAR*-like backgrounds is that the background peaks selected for motif-containing peaks tend to contain the motif, thus potentially obfuscating the motif signal. To avoid this, we include the option of downscaling the relative importance of background peaks containing the foreground motif using a parameter *λ* (disabled by default, can be set via the l argument). We track the network overlap by computing **S** = **WA**^*T*^, where an element *S*_*m,p*_ counts the total number of neighbors of peak *p* that harbor motif *m*. We then compute a soft-masked denominator matrix **D** for active motif-peak pairs:

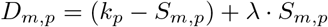

A regularized primary projection matrix 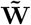 is formed by setting its non-zero entries to 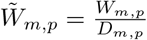. To perform the soft suppression across the adjacency graph purely through sparse matrix operations, the final corrected projection matrix **W**_cor_ is evaluated algebraically:

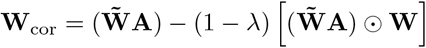

where ⊙ represents the element-wise product.

### chromVAR

*ChromVAR* (v1.32.0)^[9]^ quantifies motif accessibility by computing deviations of fragment counts across motif-containing peaks relative to matched background peaks with similar GC content and accessibility. Bias-corrected deviations are summarized as z-scores for each motif in each sample or cell. For differential analysis, we applied both the built-in `chromVAR::differentialDeviations` and *limma* v3.66.0^[28]^ to the z-score matrix (*chromVAR*.*limma*).

### Differential analysis

Unless specified otherwise (i.e. `chromVAR::differentialDeviations`), differential analysis was done using *limma* v3.66.0^[28]^. We previously showed that its moderated statistics outperformed standard *t*-test or regression analysis ^[2]^.

### Weight model

#### Fragment weights

The GC content and fragment length (FL) of sequencing fragments do not follow a uniform distribution. To avoid overrepresentation of fragments in a small number of bins when using equal-width partitioning, fragments are assigned to a two-dimensional grid *B* defined by equal-frequency bins for GC content (*bs*_GC_) and fragment length (*bs*_FL_), using user-specified numbers of divisions (arguments `nGCBins` and `nWidthBins`). Specifically, quantiles of GC content and fragment length are computed, and each fragment is assigned to a bin according to the corresponding quantile intervals. This approach ensures that each bin contains a similar proportion of fragments (Supplementary Figure S2b).

To correct for sample-specific biases, fragment-level weights are computed within each bin for each sample. These weights are defined such that the weighted fragment distribution of each sample matches the average distribution across all samples within each bin. As a result, all samples share a common distribution of weighted fragment counts across the two-dimensional GC–FL grid, effectively removing biases associated with GC content and fragment length (Supplementary Figure S2c).

#### Peak weights

A peak-level accessibility count matrix is constructed using the weighted fragments. This matrix is used to generate an MA plot and is further normalized at the peak level using affy::normalize.loess (v1.88.0)^[29]^, a loess-based normalization method.

For each pair of samples *i* and *j*, the log fold change (*M*) and average log-count (*A*) are computed for each peak. A loess curve is then fitted to *M* as a function of *A*, and the fitted trend is subtracted to correct for systematic biases. The span parameter is set to 0.3, and the loess family is specified as `symmetric`. To reduce fitting time, peaks are downsampled to 10 000 based on their mean accessibility. Peaks are assigned to 100 equal-size accessibility bins according to their average log-count, and a maximum of 100 peaks are sampled without replacement from each bin (i.e. bins with fewer peaks are neither subsampled). The stratified sampling strategy ensures that peaks at the extreme end of accessibility are taken into account in fitting the loess curve.

### Insertion model

#### Position-weighted insertion counts

For each match of motif *m* a genomic window *W* is established by extending a fixed margin *e* (default: ±200 bp, extension argument) around its center. Insertions are counted at each relative position *p* ∈ *W* (applying user specified shifts via the `shift` argument, e.g. for the Tn5-insertion offset) around each motif match. Windows corresponding to motif matches on the negative strand are reversed. To calculate per motif position-weights, the sample-wise insertion count matrices are normalized such that total coverage of each sample is of equal weight. Then a consensus weight profile is computed by first summing the normalized insertion counts at each relative position *p* across the motif matches of motif *m* and second averaging this aggregated signal at each position across all samples. In positions within the motif matches, which share highly similar sequence across matches for the same motif, most of the ATAC variations stems from Tn5 bias. For this reason, the insertion weights at positions within a motif match are replaced by the median weight across these positions. Subsequently, the raw insertion weights are smoothed across positions using LOWESS regression by default (argument: `smooth.span`). Smoothing effects at positions within the motif matches are cleared using the mean of the smoothed weights and the median of the unsmoothed signal. Profiles are further adjusted by subtracting a baseline profile, i.e. an average insertion profile across TFs, to account for noisy profiles of TFs without a clear footprint. The final insertion weights are scaled so that the mean weight across *p* ∈ *W* is 1 (Supplementary Figure S6). Optionally profiles can be made symmetric (argument: `symm`).

Weighted counts are obtained by multiplying the insertions at each position around the matches of a motif by the motif’s weight profile. For a given motif, only the match with the highest (weighted) counts per peak is retained, and the counts are summed across positions and peaks to generate a matrix of weighted insertion counts across all parsed motifs and samples.

#### Computing deviations

In order to compute bias-corrected deviations, a background is necessary, and while the standard peak counts can be used for this purpose, the use of the motif-specific weights creates a discrepancy between the scale of the foreground and background for each motif. To address this, we compute a scaling factor that normalizes the per-motif mean unweighted count to its mean weighted count, and use it to adjust both the background expectation and its variance. Otherwise the deviations are computed as described for `betterChromVAR`.

### Motif interaction analysis

#### Normalization across set sizes

Both the deviations and the z-scores show a correlation with the size of the set (i.e. the number of motif matches). The computation of the deviations uses the expected counts as a denominator, thus leading to smaller deviations for motifs that have many sites (and hence higher total count). Similarly, the background variability is greater for motifs that have fewer sites, and hence the z-scores are smaller. While this is desirable to identify more robust signals, these patterns of dependence are suboptimal for comparing the magnitude of changes, as one does in interaction analysis. To this end, we therefore normalize the z-scores before performing comparisons. Specifically, the bait motif’s Z-scores are scaled to their expected magnitude if the bait had as many motif matches as the interaction has (i.e. as the intersection of the two), using the relationship:

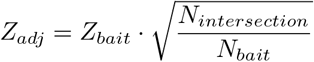

This can be achieved with the `normalizeDevsForSize` function of *betterChromVAR*.

#### Discovery of interactions

To identify motifs interacting with a given bait motif, we use linear regression via `limma`. For each motif, we fit a model on the (size-normalized) bait deviations (as baseline) and the deviations of the intersection between the motifs. We then fit a model of the form:

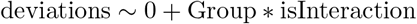

We then formulate the following contrast to capture a potential interaction (i.e. synergistic or antagonistic) effect:

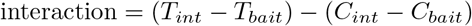

where *T* and *C* respectively represent the treatment and control groups.

Finally, variances were moderated using limma’s empirical Bayes pipeline, using the square-root of the number of intersecting matches as trend covariate.

#### Distance-dependent interaction profiling

Finally, we evaluate the candidate in more detail using the `exploreMotifInteraction` function. The function constructs an annotation by splitting sites with both motifs into equally-frequent bins based on the distance between the motifs, and compares those to ‘singlet’ sites (peaks containing only one of the two motifs). In this way, we can distinguish between direct interactions (occurring only at short distances) and broader patterns of co-occupancy persisting across long distances.

#### CVnorm

For the CVnorm bias normalization method, the matrices of samples’ counts per bias (i.e. background) bins *B* and bin-bin selection probabilities *P* are obtained as for *betterChromVAR*. Let *ϵ*_*i*_ be the global expected fraction of reads in bin *i* (e.g. calculated as the mean across cells or groups). We then calculate the smoothed observed proportions (*O*) and the smoothed background expectation (*E*) in bin *i* for each sample *s* as follows:

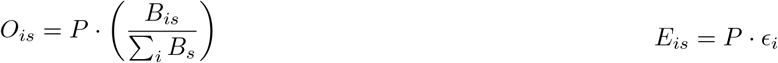

The ratio *R*_*is*_ = *O*_*is*_*/E*_*is*_ represents the raw technical bias factor for bin *i* in sample *s*. When not using variance-based shrinkage of the bias (see next section), the corrected counts for peak *j* (from bin *i*) in sample *s* can then be obtained as:

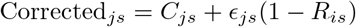

Where *C*_*js*_ is the original counts, and *ϵ*_*js*_ is the expected counts (i.e. the proportion of reads in peak *j* in the average across samples, multiplied by total read count in sample *s*). Eventual negative values are set to zero, and by default, original zeros are preserved as such.

#### Variance-based bias shrinkage

To avoid the normalization ‘correcting away’ large biological differences between experimental groups, we implement an optional weighting scheme similar to that used by smooth quantile normalization method (see the *qsmooth* package) ^[22]^. Briefly, we first log-transform the bias to make it symmetrical, and then compute, for each bias bin *i*, the sum of square differences to the mean log-transformed bias of bin *i* across samples of the same group (*SSW*), as well as the between groups sum of squares *SSB*. The mean squares, accounting for the degrees of freedom, is then:

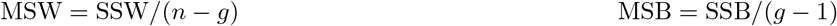

Where *n* is the number of samples and *g* the number of groups.

From this, we can then compute a weight *w*_*i*_ for each bin *i* is as

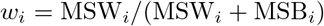

By default, the log-transformed bias is multiplied by this weight before applying the correction. In this way, if the bias varies a lot between replicates, *w* approaches 1, and we apply the full correction. If instead if varies most between groups, *w* approaches 0 and little or no correction is applied to peaks of that bin. Note that this is different from the approach used in *qsmooth*: in the latter, when variance is explained by groups, the between-group differences are not corrected for, but the within-group differences are. While such a behavior can be turned on in `CVnorm`, it leads to increased False Discovery Rate (FDR) in downstream differential analysis, and is therefore disabled by default.

## Availability of data and materials

The Cusanovich dataset is available at Gene Expression Omnibus (GSE174280). The fragment files for the SpearATAC-seq datasets (GSE168851: GATA1, GATA2, RUNX1, RUNX2, KLF1, and MYC) were downloaded from GEO. Raw sequencing reads for the other datasets (NR1H3/4: GSE149075; BANP: GSE155604) were downloaded from SRA.

The *epiwraps* package is available at https://github.com/ETHZ-INS/epiwraps and has been submitted to Bioconductor. The *betterChromVAR* package is available at https://github.com/plger/betterChromVAR and on Bioconductor. The *weightedMotifAccess* package is available at https://github.com/Jiayi-Wang-Joey/weightedMotifAccess and will be submitted to Bioconductor.

The code to reproduce the analyses and figures is available at https://github.com/ETHZ-INS/motif_analysis_paper.

## Backmatter

## Acknowledgments

The authors wish to thank Michael Stadler for feedback and discussion.

## Authors’ contributions

**Jiayi Wang** Formal analysis, Investigation, Methodology, Software (weight/insertion models), Writing – original draft

**Emanuel Sonder** Conceptualization, Formal analysis, Methodology, Software (weight/insertion models), Writing – original draft

**Silvia Domcke** Conceptualization (interactions), Writing – Review & Editing

**Mark D. Robinson** Supervision, Funding acquisition, Writing – Review & Editing

**Pierre-Luc Germain** Conceptualization, Formal analysis, Investigation, Methodology, Software (betterChromVAR, epiwraps), Funding acquisition, Supervision, Writing – original draft

## Supplementary Figures

**Supplementary Figure S1:**
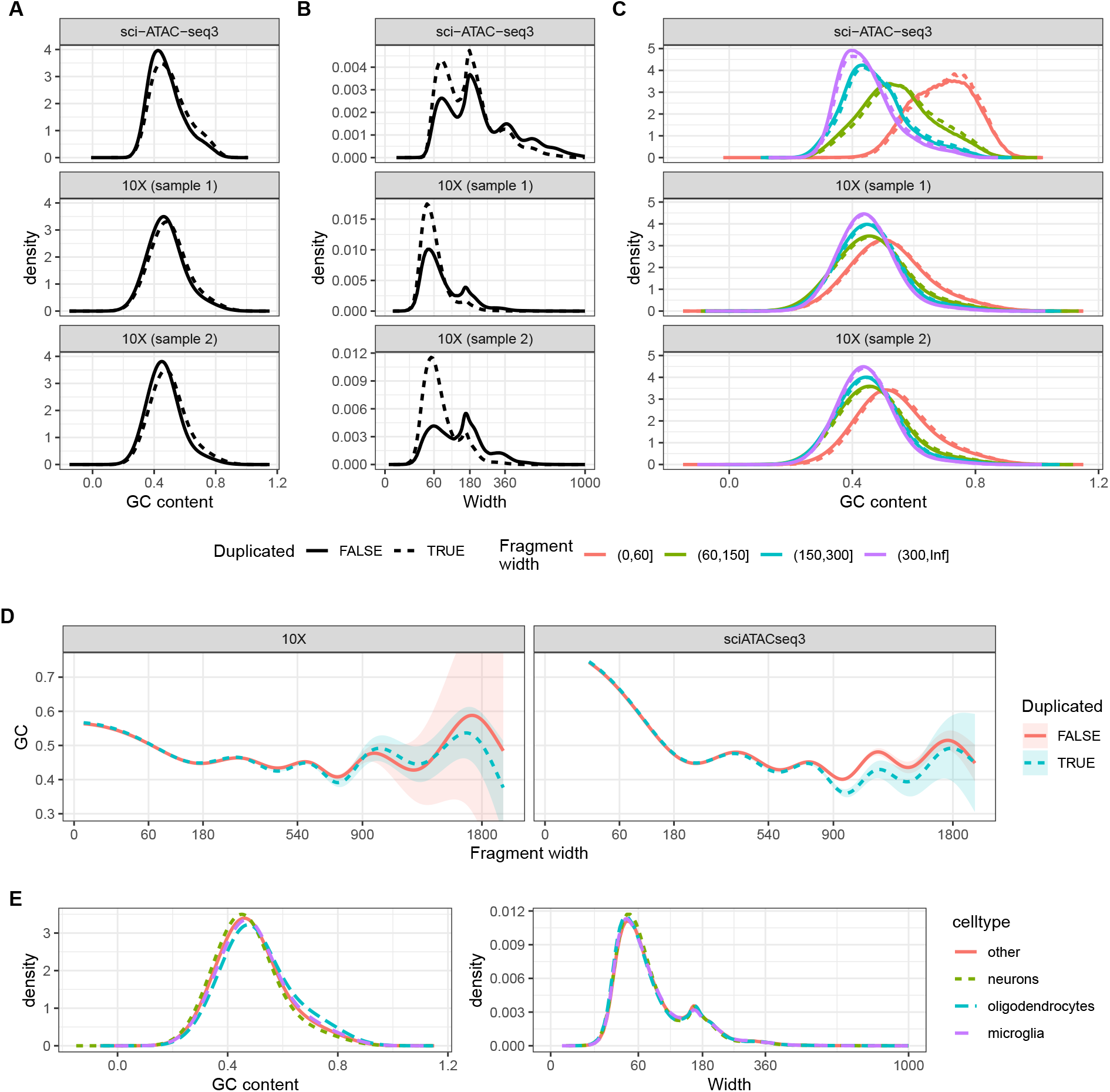
Exploration of PCR bias. using two single-cell ATAC-seq samples generated using 10X ^[19]^, as well as a sample generated by sci-ATAC-seq3 ^[30]^. **A:** Duplicated fragments tend to have a slightly higher GC content. **B:** Duplicated fragments tend to be much shorter. **C-D:** The GC bias in duplicated fragments is chiefly attributable to PCR’s bias towards shorter fragments. **D** represents a generalized additive model (*gc ~ s*(*width*)) for each type of fragment, showing no amplification bias except possibly with large (and very low-abundance) fragments. The oscillation pattern roughly antiphase with the fragment length distribution, most likely attributable to ‘hybrid’ fragments containing both nucleosome-containing and -free regions that increase the latter’s GC content. **E:** Different cell types within the same single-cell ATAC-seq sample (sample 1 above from ^[19]^) show no difference in fragment length distribution, and very limited differences in GC content.

**Supplementary Figure S2:**
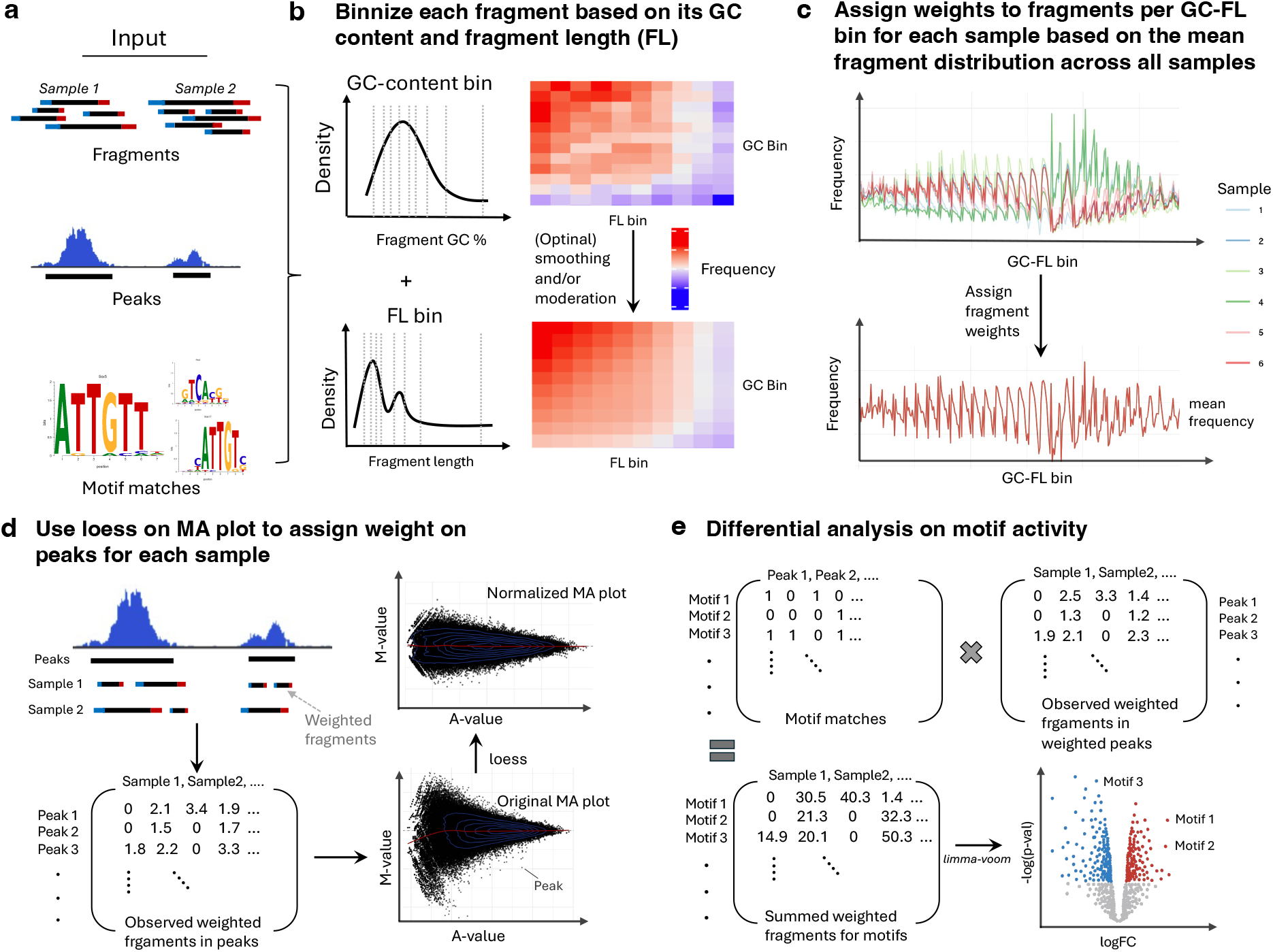
Weight model workflow. **A:** The weight model takes ATAC-seq fragments, peaks, and motif matches matrix as input for each sample. **B:** Each fragment is assigned to a GC-FL bin based on its GC-content or fragment length using quantile binning, with optional smoothing or moderation. **C:** A fragment weight is assigned to each GC-FL bin in each sample to adjust sample-specific fragment frequency distributions to match the sample-mean distribution. **D:** The weighted fragments are summed for each peak, resulting in a weighted peak-level accessibility count matrix. Cyclic loess normalization is applied to assign peak weights: a loess curve is fitted to the MA plot of each distinct pair of samples, where the M-value on the y-axis is logFC in ATAC signal between two samples and the A-value on the x-axis is the average accessibility across those two samples. **E:** The motif matches matrix is multiplied by the peak accessibility count matrix to generate a motif accessibility matrix. Next, we transformed the motif accessibility matrix using `voom` and identify differentially-active motifs using `limma`.

**Supplementary Figure S3:**
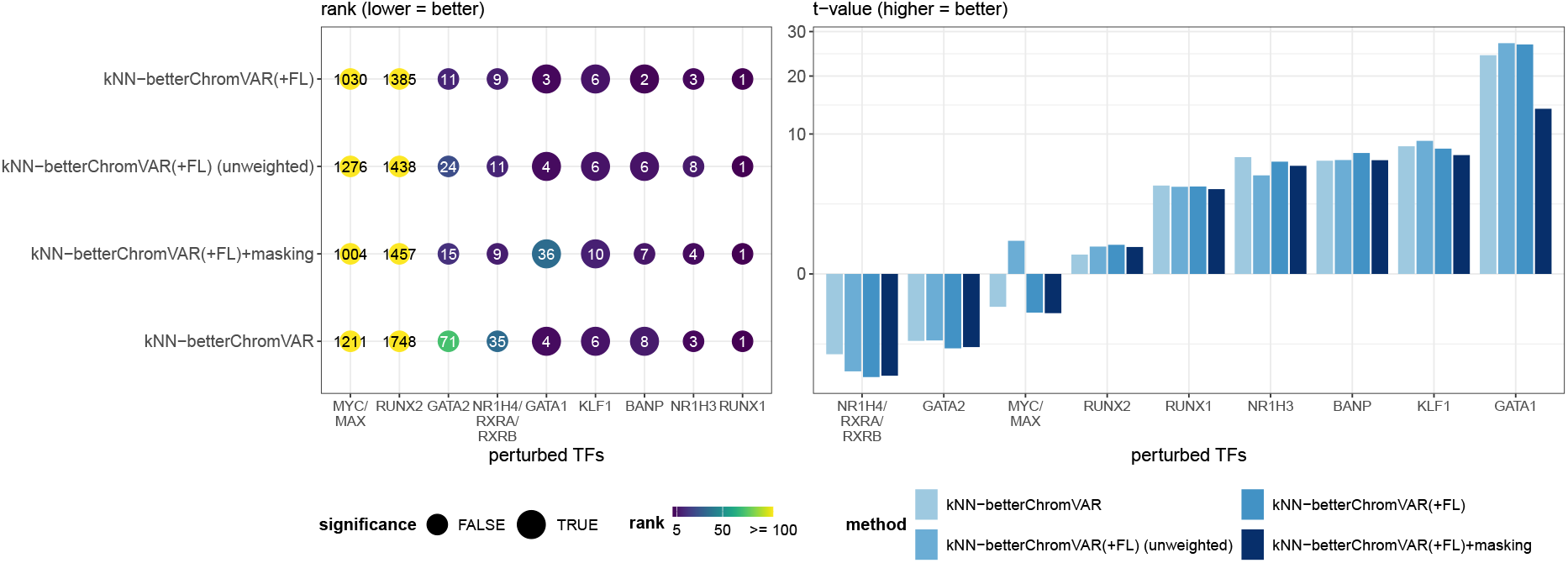
Same benchmark as shown in Figure 2, but including the different kNN-based variants of betterChromVAR.

**Supplementary Figure S4:**
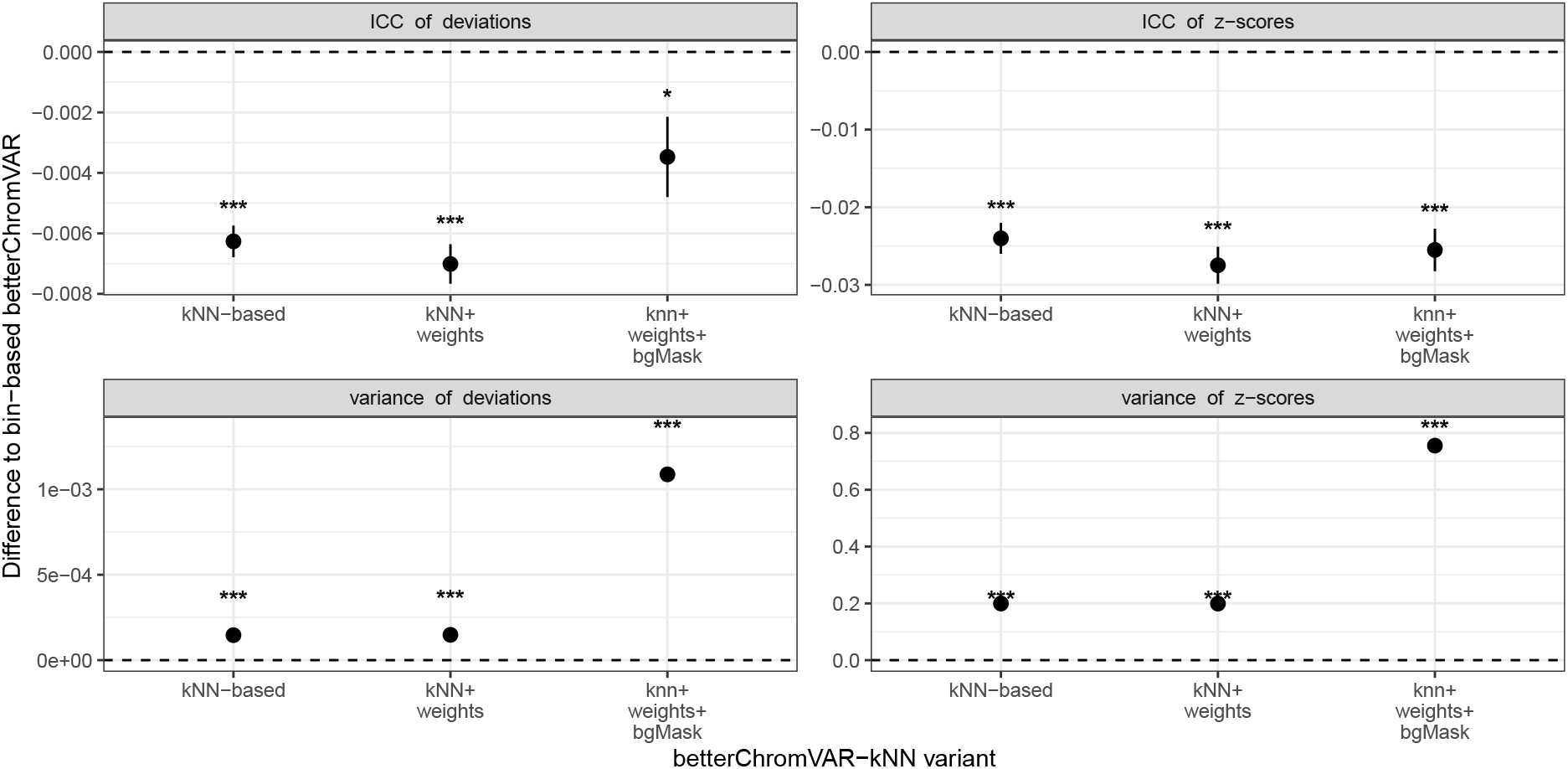
Comparison of the kNN-based *betterChromVAR* variants to bin-based *betterChromVAR*. Shown is the difference in mean intra-class correlation (ICC, top row) or mean motif variance (bottom row) between each kNN-based variant and the bin-based *betterChromVAR*, using data from Pierce *et al*. [31]. Each is computed comparing cells harboring the same guide RNA (and hence expected to be comparable), either using the motif deviations (left) or z-scores (right), and differences to bin-based are computed on the means. For computational efficiency, ICC is approximated as the correlation of each cell to the mean of the cells having the same gRNA. ‘kNN+weights’ refers to the dimensional weighting before computing kNN, and ‘+bgMask’ refers to additionally masking background peaks harboring the foreground motif. Shown are the mean and standard error across gRNAs; stars represent the significance of a one-sample *t*-test against an expectation of zero difference: *= p < 0.05, **= p <0.01, and ***= p*<*0.001.

**Supplementary Figure S5:**
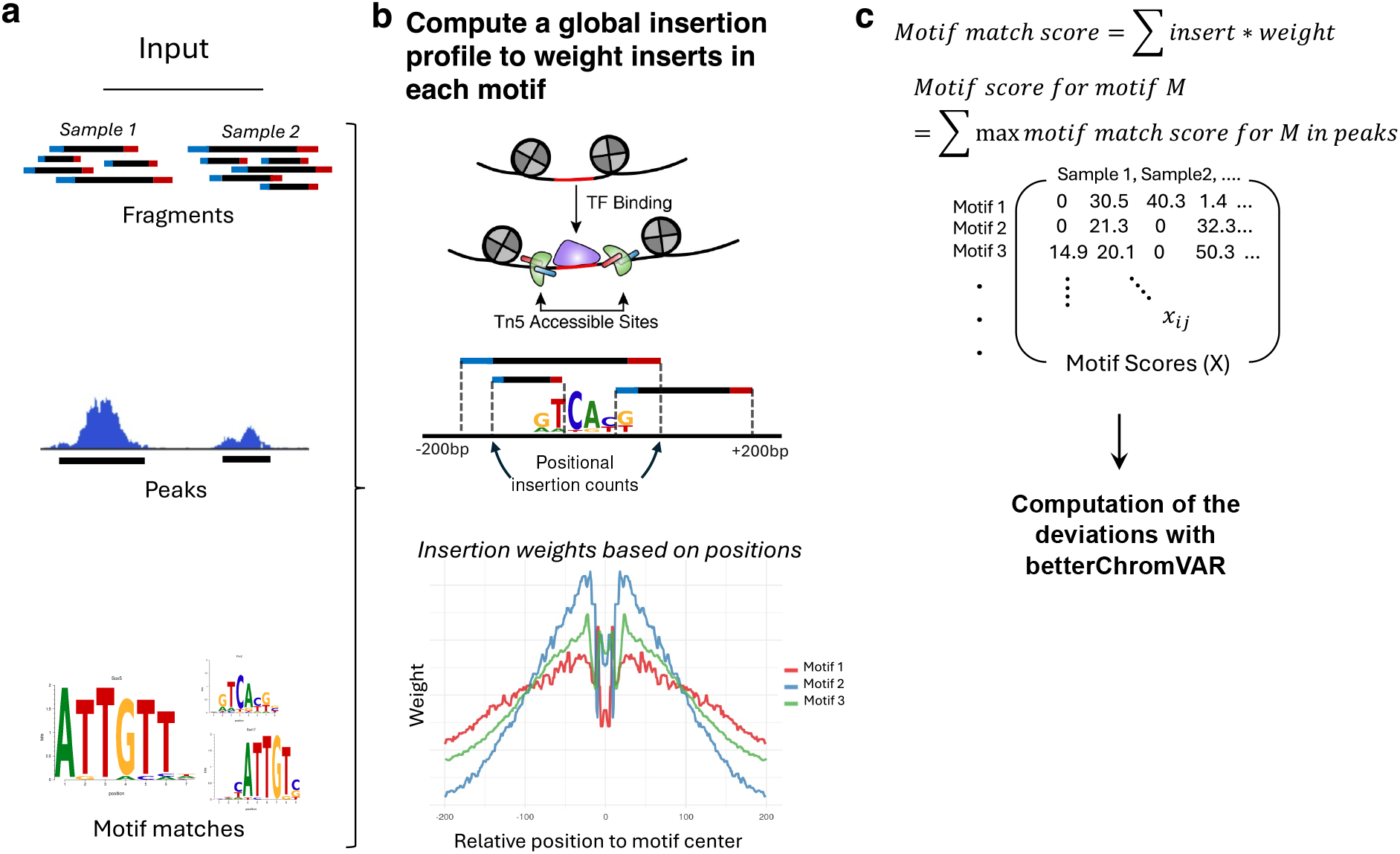
Insertion model workflow. (a) The insertion model takes ATAC-seq reads, peaks and motif match positions as input. (b) The binding of a transcription factor (TF) prevents Tn5 insertions and leaves a footprint. A global insertion profile is computed for each motif to assign weights to each insertion based on its position relative to the motif center in the flanking region. (c) The score for a specific motif match is simply the sum of the weighted insertions. The motif score for a motif *M* is computed as the sum of its maximum motif match scores across all peaks. Motif deviations are then computed using *betterChromVAR*.

**Supplementary Figure S6:**
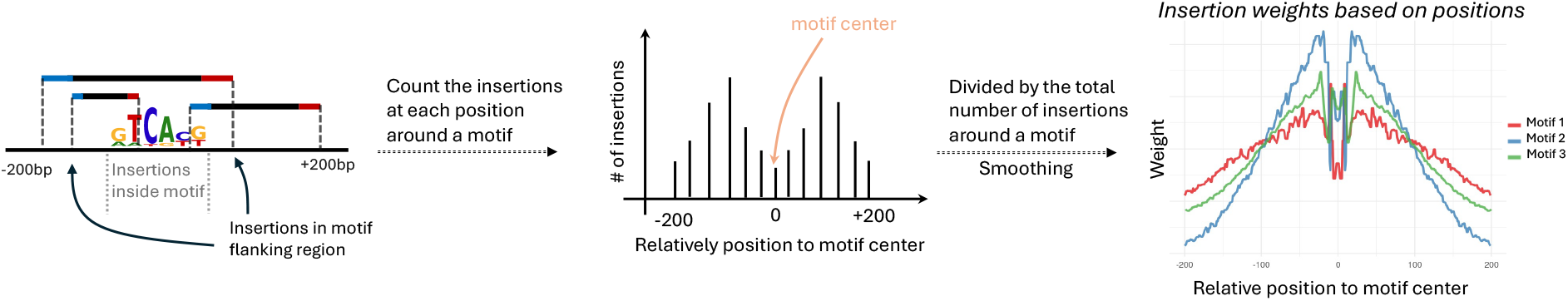
Calculation of a global insertion profile for each motif.

